# Multimodal, multiscale connectivity blueprints of the cerebral cortex

**DOI:** 10.1101/2022.12.02.518906

**Authors:** Justine Y. Hansen, Golia Shafiei, Katharina Voigt, Emma X. Liang, Sylvia M. L. Cox, Marco Leyton, Sharna D. Jamadar, Bratislav Misic

## Abstract

The brain is composed of disparate neural populations that communicate and interact with one another. Although fiber bundles, similarities in molecular architecture, and synchronized neural activity all represent brain connectivity, a comprehensive study of how all these connectivity modes jointly reflect brain structure and function remains missing. Here we systematically integrate seven multimodal, multiscale brain connectivity profiles derived from gene expression, neurotransmitter receptor density, cellular morphology, glucose metabolism, haemodynamic activity, and electrophysiology. We uncover a compact set of universal organizational principles through which brain geometry and neuroanatomy shape emergent connectivity modes. Connectivity modes also exhibit unique and diverse connection patterns, hub profiles, dominant gradients, and modular organization. Throughout, we observe a consistent primacy of molecular connectivity modes—namely correlated gene expression and receptor similarity—that map well onto multiple phenomena including the rich club and patterns of cortical abnormalities across 13 neurological, psychiatric, and neurodevelopmental disorders. Finally, we fuse all seven connectivity modes into a single multimodal network and show that it maps onto major organizational features of the brain including structural conenctivity, intrinsic functional networks, and cytoarchitectonic classes. Altogether, this work contributes to next-generation connectomics and the integrative study of inter-regional relationships.

## INTRODUCTION

Brain connectivity classically refers to the physical neural fibers that link disparate neuronal populations. Axonal projections can be reconstructed by imaging fluorescently-labelled proteins that are either injected into or genetically expressed by a cell, or by stacking electron microscopy images of thinly sliced brain sections [78, 82]. At the macroscale, diffusion-weighted imaging can be used to trace large fiber bundles that connect pairs of brain regions *in vivo*, which collectively constitute the structural connectome [54, 136]. Across organisms, spatial scales, and imaging techniques, the brain’s structural architecture presents hallmark features including a prevalence of short range connections resulting in functionally segregated modules [134], and a small number of disproportionately densely interconnected hubs [145]. Ultimately, studying the brain’s structural connectome has advanced our understanding of how information is transmitted [8, 124], how brain structure supports function [137], and how perturbations may result in network-defined pathology spread [161].

However, the structural network alone abstracts away the molecular and physiological heterogeneity that exist in the brain. An alternative representation of connectivity is feature similarity: if two brain regions exhibit similar features, we might expect them to be related to one another, or “connected” [1]. This approach is already widely used on the haemodynamic BOLD signal but also exists for time-series measures from other imaging modalities such as magneto-/electroencephalography (MEG/EEG) and dynamic FDG-fPET (all called “functional connectivity”) [24, 32, 46, 52, 70]. In cases where multiple measures of a feature exist at each brain region, such as gene expression levels across many genes, connectivity can represent the similarity of brain regions with respect to a single local feature [56, 57, 60, 103, 111, 122, 125]. In each case, the ensuing region × region network represents a form of connectivity between brain regions.

As multiple layers of connectivity, or “connectivity modes”, become available, it becomes possible to integrate them into a single framework and deduce how these layers interact with one another, and in what ways they are unique or complementary. For example, brain structure is heterogeneously coupled to haemodynamic functional connectivity along the sensory-association cortical hierarchy [14, 150]. Information about inter-regional feature similarity adds additional insight on how structure supports function, and has been shown to improve the structure-function concordance [56, 91, 105]. The advance in neuroimaging techniques and data sharing standards has now made it possible to study multiple connectivity modes jointly, spanning a range of spatial and temporal scales. The future of connectomics is therefore no longer limited to structural connectivity, but can be approached from a multi-modal, multi-scale angle.

Here we integrate seven layers of connectivity, including gene expression, receptor density, cellular composition, metabolism, and electrophysiology, to assemble a comprehensive, multiscale wiring blueprint of the cerebral cortex. First, we establish the common and unique manners in which connectivity modes reflect brain structure and geometry. Next, we identify cross-modal hubs and consistently high-strength edges. We then test how different connectivity modes capture patterns of cortical abnormalities across 13 neurological, psychiatric, and neurodevelopmental disorders. Moreover, we show that connectivity modes demonstrate diverse gradient and modular decompositions. Finally, we iteratively fuse all seven connectivity modes into a single harmonized network. All seven connectivity profiles are publicly available in three parcellation resolutions (https://github.com/netneurolab/hansen_many_networks), in hopes of facilitating integrative, multi-scale analysis of human brain connectivity.

## RESULTS

For each brain feature, a similarity network can be represented as a region × region matrix. Rows and columns represent brain regions, and elements—the edges of the similarity network—represent how similarly two brain regions present the specific feature. This similarity can also be thought of as connectedness, such that two brain regions that share similar features are considered strongly connected. A comprehensive assessment of cortical connectivity modes was achieved by constructing and analyzing seven different connectivity matrices, spanning multiple spatial and temporal scales. These include: (1) correlated gene expression, describing transcriptional similarity across *>* 8 000 genes from the Allen Human Brain Atlas [59]; (2) receptor similarity, describing how correlated pairs of brain regions are in terms of protein density of 18 neurotransmitter receptors/transporters [56]; (3) laminar similarity, describing how correlated pairs of brain regions are in terms of cell-staining intensity profiles from the Big-Brain atlas [4, 103]; (4) metabolic connectivity, measured as the correlation of dynamic FDG-PET (i.e. glucose uptake) time-series [69, 70]; (5) haemodynamic connectivity, measured as the correlation of fMRI BOLD time-series from the Human Connectome Project (HCP) [147]; (6) electrophysiological connectivity, measured as the first principal component of MEG connectivity across six canonical frequency bands from the HCP [126, 147]; and (7) temporal profile similarity, describing how similar pairs of brain regions are in terms of temporal features of the fMRI BOLD signal [128]. To facilitate comparison between networks, each network was parcellated to 400 cortical regions and edge values underwent Fisher’s *r*-to-*z* transform [120]. Networks were also parcellated to 100 and 68 cortical regions for the sensitivity and replication analyses (see *Sensitivity and replication analysis*).

### Common organizational principles of connectivity modes

In Fig. 1a, we visualize each normalized connectivity matrix as a heatmap where the colourbar limits are − 3 and 3 standard deviations of the edge weight distribution (for edge weight distributions, see Fig. S1a). Brain regions are ordered by left then right hemisphere. Within each hemisphere, regions are further stratified by their membership in the seven canonical intrinsic functional networks (Schaefer-400 parcellation [120, 159]). Homotopic connections stand out, indicating that homologous brain regions in left and right hemispheres are consistently similar to each other (Fig. S1b). Visually, each connectivity mode demonstrates non-random network organization, which we explore in subsequent sections. Furthermore, similarity between brain regions decreases as the distance between brain regions increases (Fig. 1b), consistent with the notion that proximal neural elements are more similar to one another [45, 56, 62, 111, 128]. In all cases except laminar similarity, this relationship is better fit by an exponential rather than linear function.

**Figure 1.**
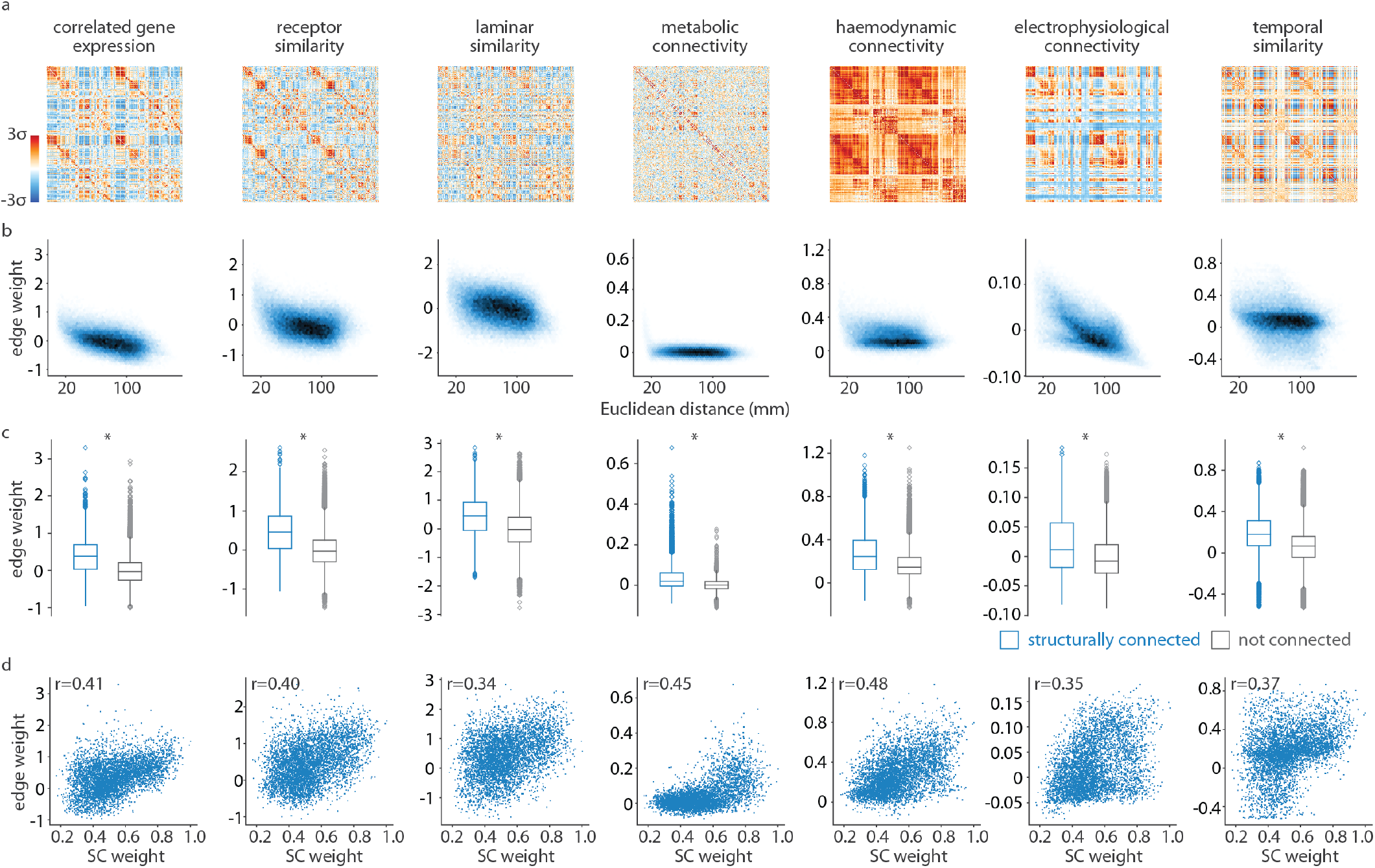
Common organizational principles of connectivity modes. Each connectivity mode is a normalized similarity matrix, where elements of the matrix index how similarly two brain regions present a specific feature. (a) Connectivity modes are shown as heatmaps, ordered according to the 400-region Schaefer parcellation [120]. The colourbar limits are − 3 to 3 standard deviations of the edge weight distribution. (b) Edge weights between every pair of brain regions (i.e. upper triangular elements) decrease with Euclidean distance across all seven connectivity modes. Darker colour represents greater density of points. (c) Edge weight distributions are visualized separately for edges that also exist in the structural connectome (blue) and those that do not (grey), according to a group-consensus structural connectome from the HCP [147]. Structurally connected brain regions show greater similarity than regions that are not structurally connected. Boxplots represent the 1st, 2nd (median) and 3rd quartiles, whiskers represent the non-outlier end-points of the distribution, and diamonds represent outliers (*>* 1.5 inter-quartile range). (d) For edges that also exist in the structural connectome, connectivity mode edge weight increases with the strength of the structural connection.s

We first sought to relate each connectivity mode to the brain’s underlying structural architecture. We constructed a weighted structural connectome using diffusion-weighted MRI data from the Human Connectome Project; this network represents whether, and how strongly, two brain regions are connected by white matter streamlines. We find that, across all modalities, brain regions that are physically connected by white matter show greater similarity than those that are not connected (Fig. 1c). These differences are greater than in a population of degree- and edge length-preserving surrogate structural connectomes, suggesting that the effect is specifically due to wiring rather than spatial proximity [20]. Finally, for brain regions that are structurally connected, we find a correlation between the strength of the structural connection and the similarity between brain regions, regardless of brain feature (Fig. 1d) [65]. Altogether, we find that connectivity modes demonstrate common organizational principles that respect geometry, neuroanatomy, and anatomical connectivity.

### Structural and geometric features of connectivity modes

Although connectivity modes share organizational properties, the median correlation between networks is *r* = 0.25 (range: *r* = 0.10–0.53; Fig. S2). In other words, networks are not redundant and we would expect each level of organization to make a unique contribution to brain topography. To directly compare edge weights across networks, we converted edge weights to ranks, such that the smallest (i.e. most negative) edge is ranked 1 and the strongest (i.e. most positive) edge is ranked 79 800 (equal to the number of edges in each network). We focus on two metrics to classify edges between brain regions: distance and structural connectivity.

First we consider whether some brain features show more dominant edge strength for long-range communication than others. Two brain regions that exist at spatially distant locations may communicate via a structural connection but the long distance would result in signal attenuation and a conduction delay. However, if both brain regions exhibit similar features, their activity might be more easily synchronized irrespective of distance. To explore this possibility, we bin all 79 800 edges into fifty equally-sized bins (1 596 edges per bin). For each connectivity mode separately, we calculate the median edge rank within each bin (Fig. 2a). Median edge rank decreases as the distance between brain regions in each bin increases, consistent with our finding in Fig. 1b. The brain features that retain the strongest edge strength when regions are distant are receptor similarity, temporal similarity, haemodynamic connectivity, and metabolic connectivity, in order of median strength within the last (longest-distance) bin. On the other hand, electrophysiological connectivity shows especially low long-distance edge rank and high short-distance edge rank.

**Figure 2.**
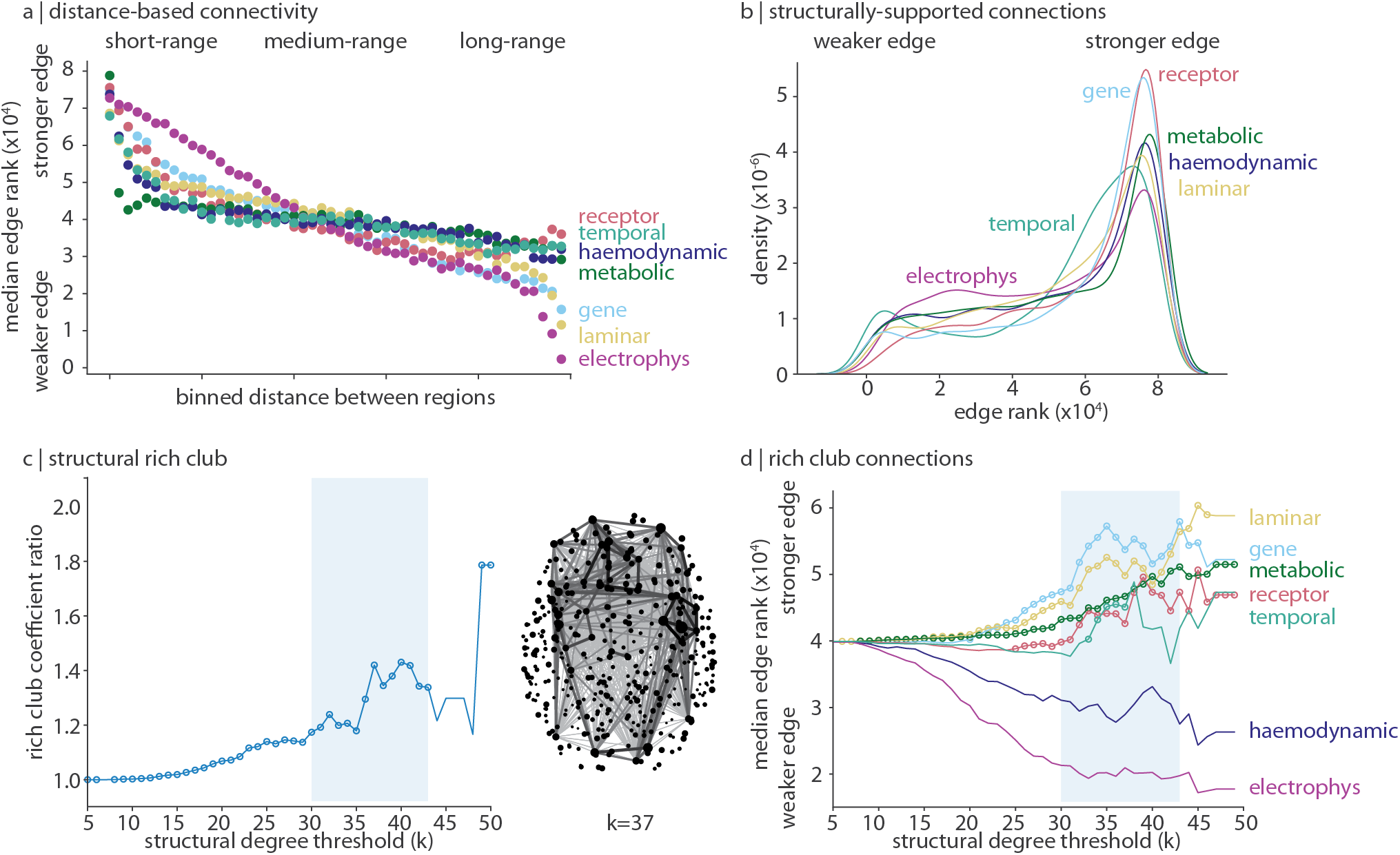
Structural and geometric features of connectivity modes. To compare edge weights across networks, edges are rank-transformed. (a) Edges are binned into 50 equally-sized bins of increasing Euclidean distance (1 596 edges per bin). For each connectivity mode, the median edge rank is plotted within each bin. (b) A kernel density estimation is applied on the edge rank distribution of edges that also have a structural connection, for each connectivity mode. (c) For a structural degree threshold *k* ∈ [5, 50], we calculate the rich club coefficient ratio and show a characteristic increase in rich club coefficient ratio when 30 ≤ *k* ≤ 43. Circles indicate structural degree thresholds where the rich club coefficient ratio is significantly greater than a null distribution of ratios calculated using a degree-preserving rewired network (1 000 repetitions). On the right we show the set of structural edges connecting regions with structural degree ≥ 37. Edge shade and thickness are proportional to edge weight, and point size are proportional to structural degree. (d) For each *k* ∈ [5, 50] and for each connectivity mode, we calculate the median edge rank of structurally-supported edges that connected regions with structural degree ≥ *k*. Circles indicate structural degree thresholds where the median rich-link edge rank of a connectivity mode is significantly greater than the edge rank of all other structurally-supported edges (Welch’s t-test, one-sided).

We next shift our focus from distance to structurally connected brain regions and ask how different brain features reflect the underlying structural architecture of the brain. Given the correlation between structural connectivity and feature similarity (Fig. 1c, d), we expect to find a left-skewed distribution of structurally-supported edge ranks. We find that this property is exaggerated for receptor similarity and correlated gene expression, indicating that regions that are structurally connected demonstrate especially similar receptor density and gene transcription (Fig. 2b). We take this one step further and track how edge strength changes depending on the structural embedding of each brain region. We focus on the brain’s “rich club”: a set of disproportionately interconnected high-degree brain regions that mediates long-range information propagation and integration [47, 145].

Specifically, for each structural degree threshold *k* ∈ [5, 50] (where structural degree is defined as the number of structural connections made by a brain region), we calculate the rich club coefficient ratio on the binary structural connectome: the tendency for brain regions of degree ≥ *k* to be preferentially connected to one another, with respect to a population of degree-preserving surrogate networks. We find that the rich club coefficient ratio is inflated at approximately 30 ≤ *k* ≤ 43 (Fig. 2c). This topological rich club regime denotes a degree range where brain regions are unexpectedly densely interconnected [34]. Next, for each connectivity mode at each *k*, we calculate the median edge rank of all structurally-supported edges that link two brain regions with degree ≥ *k* (Fig. 2d). Moreover, we ask whether within-set edge ranks (i.e. edges connecting regions with degree ≥ *k*) are statistically greater than all other edges (Welch’s one-sampled t-test).

We find that edges in the brain’s topological rich club regime are especially dominated by molecular features (e.g. laminar similarity, correlated gene expression, and receptor similarity). Haemodynamic and electrophysiological connectivity are especially weak for links between high-degree regions, and temporal similarity is unstable. Metabolic connectivity is an additional connectivity mode that demonstrates significantly increased edge strength for links between high-degree regions, suggesting that energy consumption is synchronized between structural hubs. Broadly, we find that molecular connectivity modes reflect structure and geometry more than dynamic modes.

### Cross-modal hubs

Next, we explore the edges that are consistently high-strength, and the regions that make such connections. For every connectivity mode, we show an axial view of the 0.5% strongest edges (Fig. 3a). Visually, high-strength edges vary across connectivity modes: some networks form densely interconnected cores (i.e. electrophysiological connectivity and temporal similarity), some emphasize long-range (i.e. haemodynamic connectivity) or short-range (i.e. metabolic connectivity) connections, and others appear more non-specific (i.e. correlated gene expression, receptor similarity, and laminar similarity). This variability is reflected also in the hubness profiles of each connectivity modality, where a brain region’s hubness is defined as the sum of the rank-transformed edge weights between it and all other regions (Fig. 3b).

**Figure 3.**
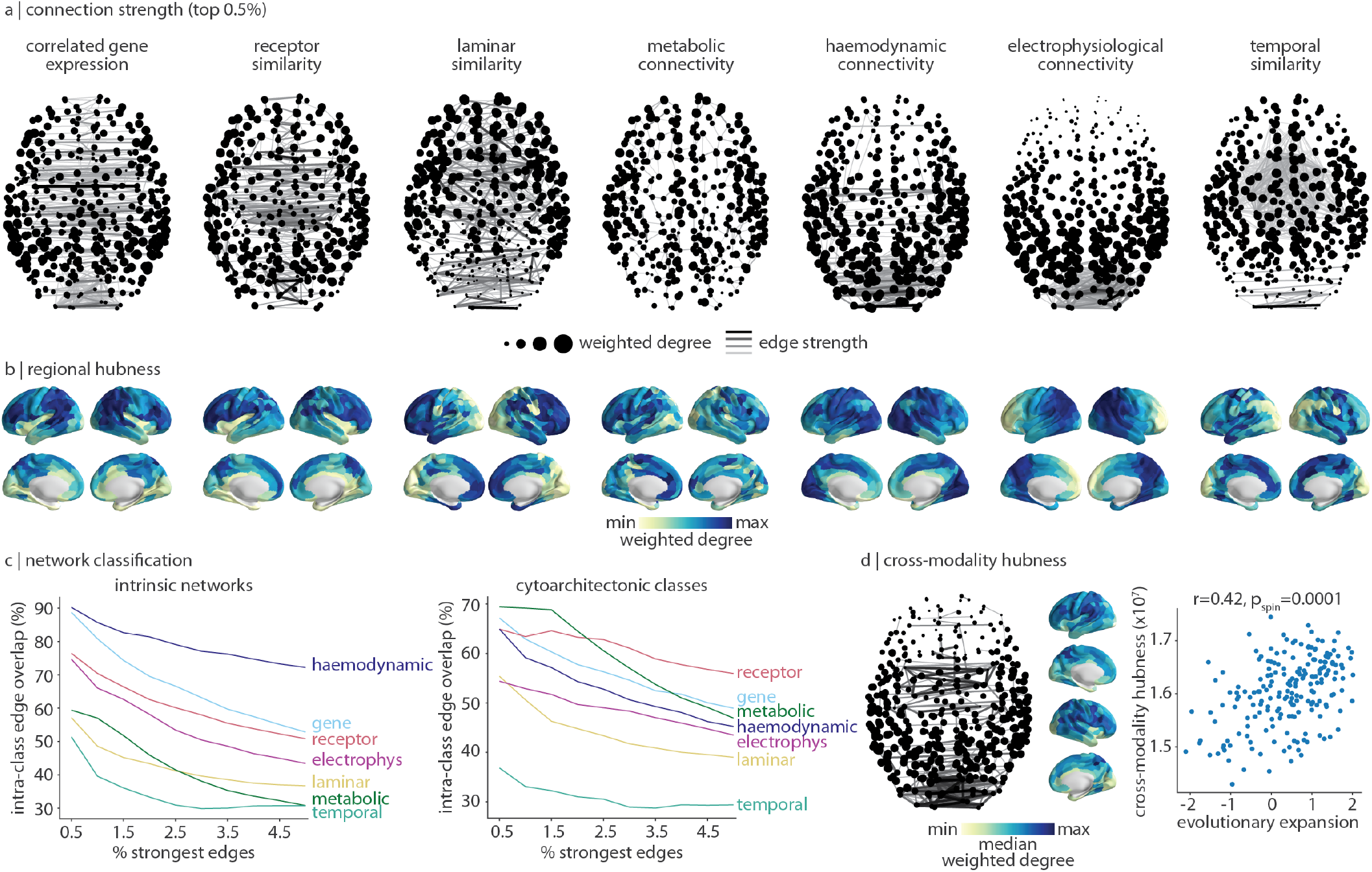
Cross-modal hubs. (a) For each connectivity mode, we plot the 0.5% strongest edges. Darker and thicker edges indicate stronger edges. Points represent brain regions and are sized according to the sum of edge weights (weighted degree). Brain views are axial, with anterior regions at the top of the page. (b) For each connectivity mode, regional hubness is defined as the sum of rank-transformed edge weights across regions. (c) For a varying threshold of strongest edges (0.5%–5% in 0.5% intervals), we calculate the proportion of edges that connect two regions within the same intrinsic network (left [159]) and cytoarchitectonic class (right [153]). (d) Across all seven connectivity modes, we calculate the median edge rank of each edge and plot the 0.5% strongest edges (left). Likewise, we calculate the median hubness (shown in panel b), which we find is significantly correlated with evolutionary cortical expansion (*r* = 0.42, *p*_spin_ = 0.0001) [63].

We were interested to know whether connectivity modalities preferentially emphasize edges that link brain regions that are functionally similar (i.e. within the same intrinsic functional networks [159]) or similar in terms of cellular composition (i.e. within the same cytoarchitectonic class [151, 153]). For a given network classification, we call edges that join two brain regions in the same network intra-class edges [125]. Next, we calculate how many of the *x* strongest edges in a given connectivity matrix overlap with intra-class edges. We let *x* vary in increments of 0.5% from 0.5% to 5% of the strongest edges in a network. We find that for every connectivity mode, the strongest edges of the network are preferentially edges that connect brain regions within the same functional and cytoarchitectonic network (Fig. 3c).

For intrinsic networks (Fig. 3c, left), the strongest edges in the haemodynamic network are almost entirely intra-class edges (90.2% for the top 0.5% strongest edges, and 72.2% for the top 5% edges). The strongest edges in correlated gene expression are also primarily intra-class edges (88.7% for the top 0.5% strongest edges) but this ratio decreases to 52.8% at 5% of the strongest edges. Meanwhile, for cytoarchitectonic classes (Fig. 3c, right), receptor similarity, correlated gene expression, and metabolic connectivity most maximize intra-class edges. Across both intrinsic and cytoarchitectonic networks, temporal similarity retains the fewest intra-class edges. Nonetheless, despite the variability in strongest edges across all seven connectivity modes, each network emphasizes edges between regions that are functionally and cytoarchitectonically similar.

Finally, we integrate the connectivity modes into a single map of high-strength edges and cross-modal hubs (Fig. 3d). We calculate the median edge rank across all networks and plot the top 0.5% edges. These edges represent connections that are consistently weighted highly across connectivity modes, and primarily connect visual, posterior parietal, and anterior temporal regions (Fig. 3d, left). Likewise, cross-modal hubness is calculated as the median hubness across connectivity modes (i.e. the median across brain plots shown in Fig. 3b). Here we find that transmodal regions such as the supramarginal gyrus, superior parietal cortex, precuneus, and dorsolateral prefrontal cortex are most consistently similar to other brain regions across all connectivity modes (Fig. 3d, right).

Why are some brain regions highly similar to many other regions across multiple spatial scales and biological mechanisms? We hypothesized that cross-modal hubs are more cognitively flexible and able to support higher-order, evolutionarily-advanced cognitive processes. We therefore correlated cross-modal hubness with a map of evolutionary cortical expansion [63]. Indeed, the identified cross-modal core coincides with brain regions that are more expanded across phylogeny (*r* = 0.43, *p*_spin_ = 0.0001). In other words, brain regions that are expanded in humans and therefore likely involved in higher-order cognition tend to communicate with many other brain regions. Ultimately, hubs that are defined using connectivity modes other than the classical structural connectome provide novel perspectives on how regions participate in neural circuits.

### Connectivity modes shape disease vulnerability

We next ask how connectivity modes shape the spatial patterning of brain disease. Emerging theories emphasize that the course and expression of multiple brain diseases is mediated by shared molecular vulnerability [57, 156]. We therefore compared connectivity modes with patterns of cortical abnormalities across thirteen different neurological, psychiatric, and neurodevelopmental diseases and disorders from the ENIGMA consortium (*N* = 21 000 patients, *N* = 26 000 controls) [57, 79, 141].

We define the “exposure” that region *i* has to region *j*’s pathology as the product between the (*i, j*)-edge strength (*c*_*ij*_) and region *j*’s abnormality (*d*_*j*_) (Fig. 4a) [31, 57, 127, 129]. Then the global disease exposure to region *i* is the mean exposure between region *i* and all other regions in the network with positive edge strength. Finally, we correlate cortical abnormality with disease exposure to determine whether the spatial patterning of the disease is informed by a connectivity mode (Fig. 4a, right). This analysis is repeated for each connectivity mode and each disorder, and correlation coefficients are visualized in Fig. 4b.

**Figure 4.**
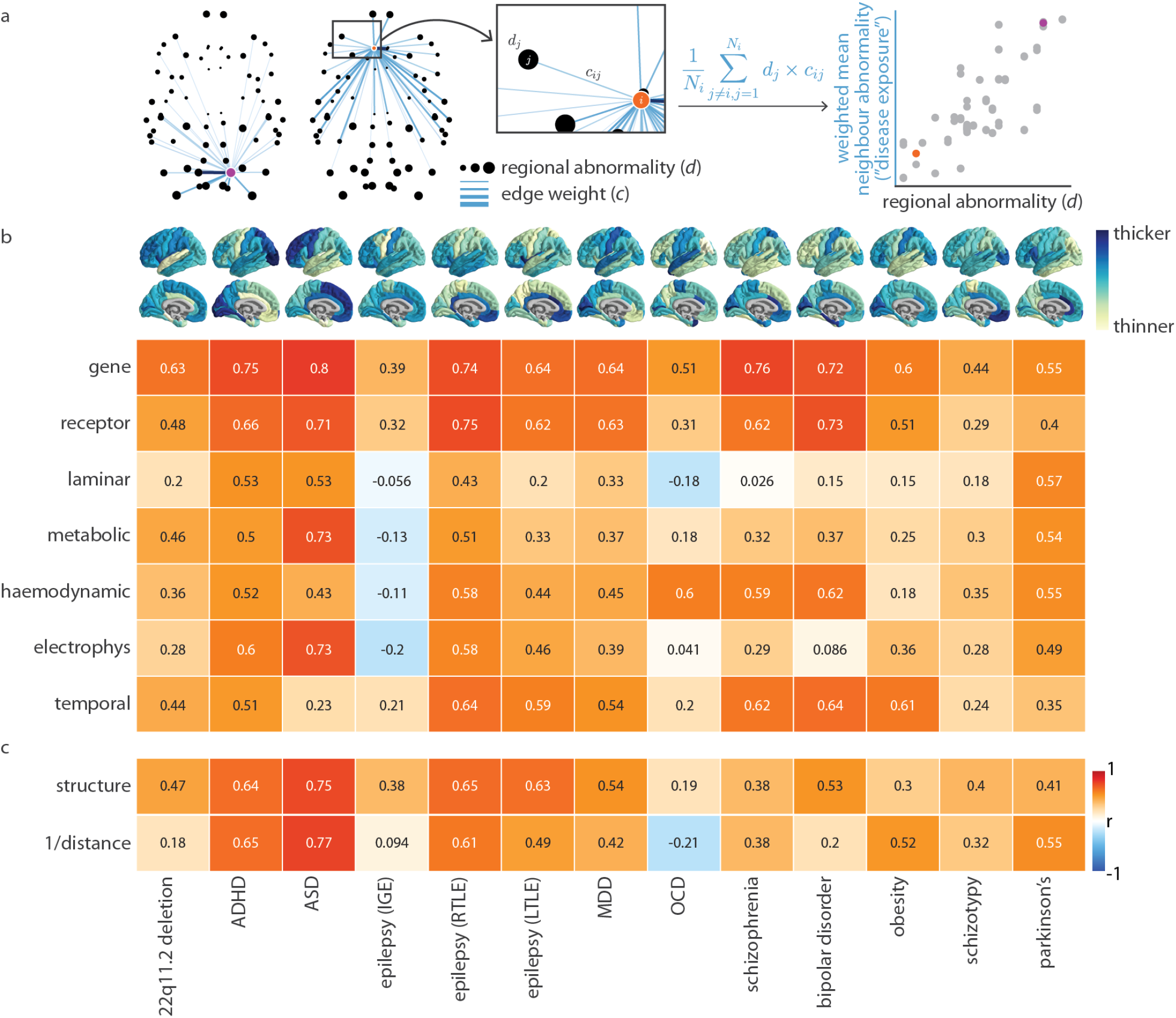
Contributions of connectivity modes to disease vulnerability. Cortical abnormality profiles for thirteen neurological, psychiatric, and neurodevelopmental disorders were collected from the ENIGMA consortium (brain plots shown in panel b; *N* = 21 000 patients, *N* = 26 000 controls [79, 141]). (a) Given a specific disorder and connectivity mode, *d*_*j*_ represents the cortical abnormality of region *j*, and *c*_*ij*_ represents the edge weight (similarity) between regions *i* and *j*. For every region *i*, we calculate the average cortical abnormality of all other regions in the network, weighted by the edge strength (“disease exposure”; note that we omit negative connections, such that *N*_*i*_ represents the number of positive connections made by region *i*). Next, we correlate disease exposure and regional abnormality across brain regions (scatter plot; points represent brain regions). We show the connectivity profiles of two example regions (highlighted in purple in the left brain network and orange in the right brain network). (b) The analytic workflow presented in panel a is repeated for each disorder and connectivity mode, and we visualize Spearman correlations in a heatmap. (c) This analysis is repeated for weighted structural connectivity (where we only consider structurally-connected regions), and Euclidean distance (where we always consider all regions in the network).

We find that correlated gene expression and receptor similarity most consistently amplify the exposure of pathology in a manner that closely resembles the cortical profile of the disease. This suggests that brain regions with similar molecular makeup may undergo similar structural changes in disease [123]. By repeating the analysis using weighted structural connectivity (in which case we only consider structurally-connected regions) and Euclidean distance between brain regions (in which case we always consider the full network), we are able to uncover cases where feature similarity amplifies disease exposure more than structure or distance alone (Fig. 4c). Cortical abnormality patterns of psychiatric disorders in particular (e.g. MDD, schizophrenia, bipolar disorder, OCD) are better explained by correlated gene expression and receptor similarity than structure or distance.

### Gradients and modules of connectivity modes

We next consider how each connectivity mode is intrinsically organized, both in terms of axes of variation (i.e. “gradients”) and network modules [37, 55, 85, 103]. We show the first principal component of each connectivity mode in Fig. 5a. Most gradients account for approximately half of the variance in the data, but the temporal similarity gradient is especially dominant (accounting for 73.8% of variance) while the metabolic connectivity gradient is especially non-dominant (accounting for 12.7% of variance; Fig. 5b). Although we might expect brain gradients to largely follow a similar sensory-association axis [68, 87, 139], we instead find that the first principal component of each connectivity mode varies considerably (median absolute correlation between gradients *r* = 0.36; Fig. 5c).

**Figure 5.**
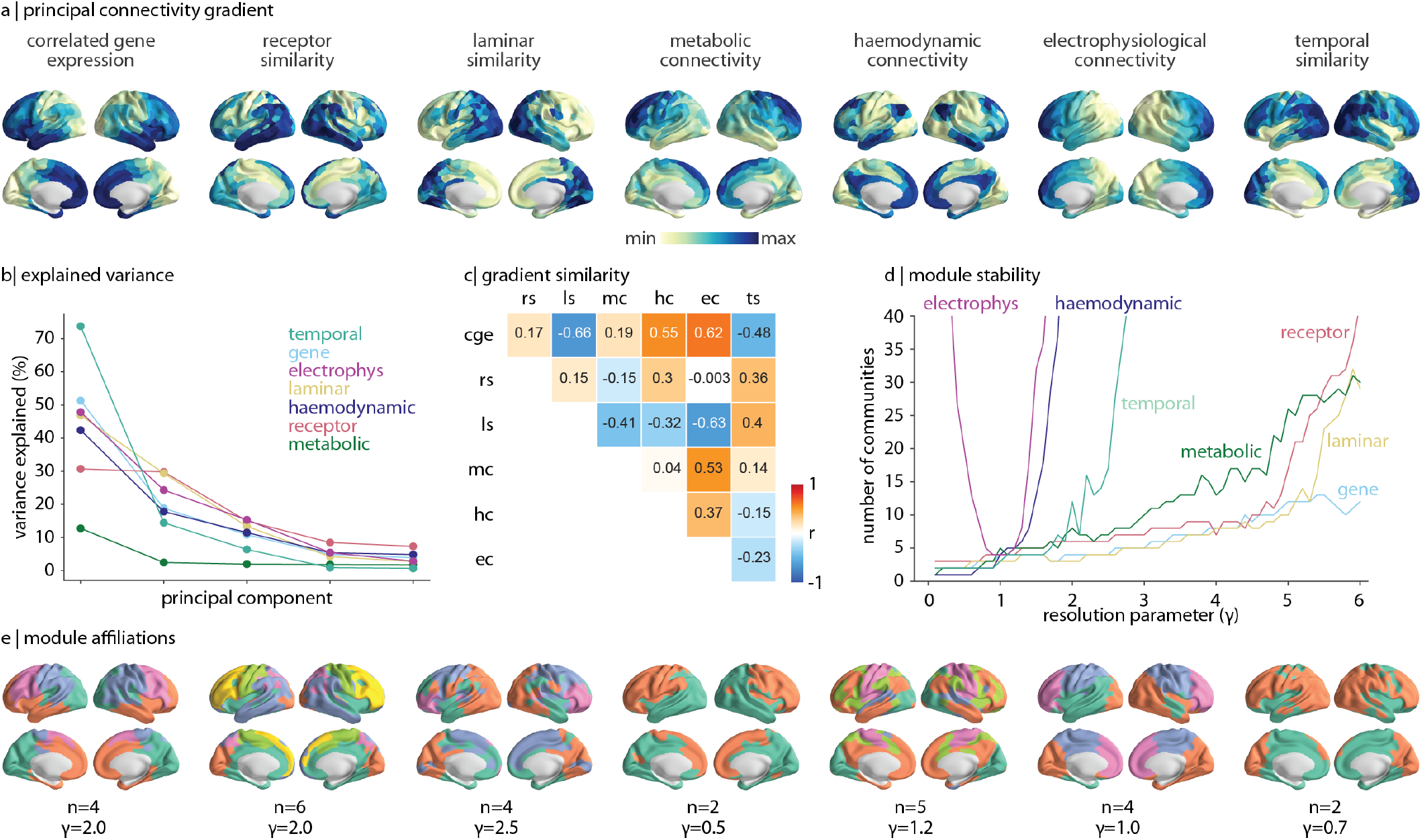
Gradients and modules of connectivity modes. (a) The first principal component (“gradient”) of each connectivity mode is shown on the cortex. (b) The percent variance explained for the first five principal components of each connectivity mode. (c) The Pearson’s correlation between every pair of network gradients, visualized as a heatmap. CGE=correlated gene expression, RS=receptor similarity, LS=laminar similarity, MC=metabolic connectivity, HC=haemodynamic connectivity, EC=electrophysiological connectivity, TS=temporal similarity. (d) The Louvain community detection algorithm is applied to each connectivity mode across different resolution parameters (0.1 ≤ *γ* ≤ 6.0, in intervals of 0.1) and the number of ensuing communities is plotted as a function of *γ*. (e) For each connectivity mode we show a single community detection solution for a specified *γ*, and we indicate the number of communities (*n*).

An alternative perspective of intrinsic network organization comes from considering whether and how the network clusters into segregated modules. We apply the Louvain community detection algorithm to each connectivity mode and, across a range of resolution parameter values (*γ*), extract community assignments for each brain region [12, 22]. To get a sense of the resolution of each network (i.e. the number of communities the network might naturally exhibit, if at all), we track the number of communities identified by the Louvain community detection algorithm across different values of *γ* (Fig. 5d). We find that the community detection solution for electrophysiology is highly unstable, with the number of identified communities changing rapidly with small changes in *γ*. The most stable solution at *γ* = 1 simply delineates the main cortical lobes. Haemodynamic connectivity and temporal similarity show a similar trend, where partitions of greater than approximately 5 networks become increasingly unstable. Meanwhile, correlated gene expression, laminar similarity, and receptor similarity show more stable community solutions, where larger changes in *γ* are required for the network to split itself into more communities. We show one possible consensus community detection solution for each network in Fig. 5e, which demonstrates that the modular organization and gradient decomposition of networks tend to be closely aligned.

### Network fusion

Ultimately, the brain is an integrated organ that coordinates each layer of connectivity. We were therefore interested in exploring a framework that merges all seven representations of brain connectivity. We rely on an unsupervised learning technique, similarity network fusion (SNF), to construct a single fused brain network [155]. SNF iteratively fuses each connectivity mode in a manner that strengthens edges that are consistently strong and weakens inconsistent (or consistently weak) edges, while giving each connectivity modality equal influence on the fusion processes. Altogether, the fused network represents a data-driven integration of each level of brain connectivity.

We find that the fused network reproduces expected relationships in the brain, including strong homotopic connections, network organization (Fig. 6a), and a negative exponential relationship with distance (Fig. 6b). In addition, structurally connected edges have significantly stronger edge weight than non-connected edges, against a degree- and edge-length preserving structural null (Fig. 6c). Finally, the fused network demonstrates a greater correlation between edge weight and structural weight (*r* = 0.53) than any of the individual connectivity modes.

**Figure 6.**
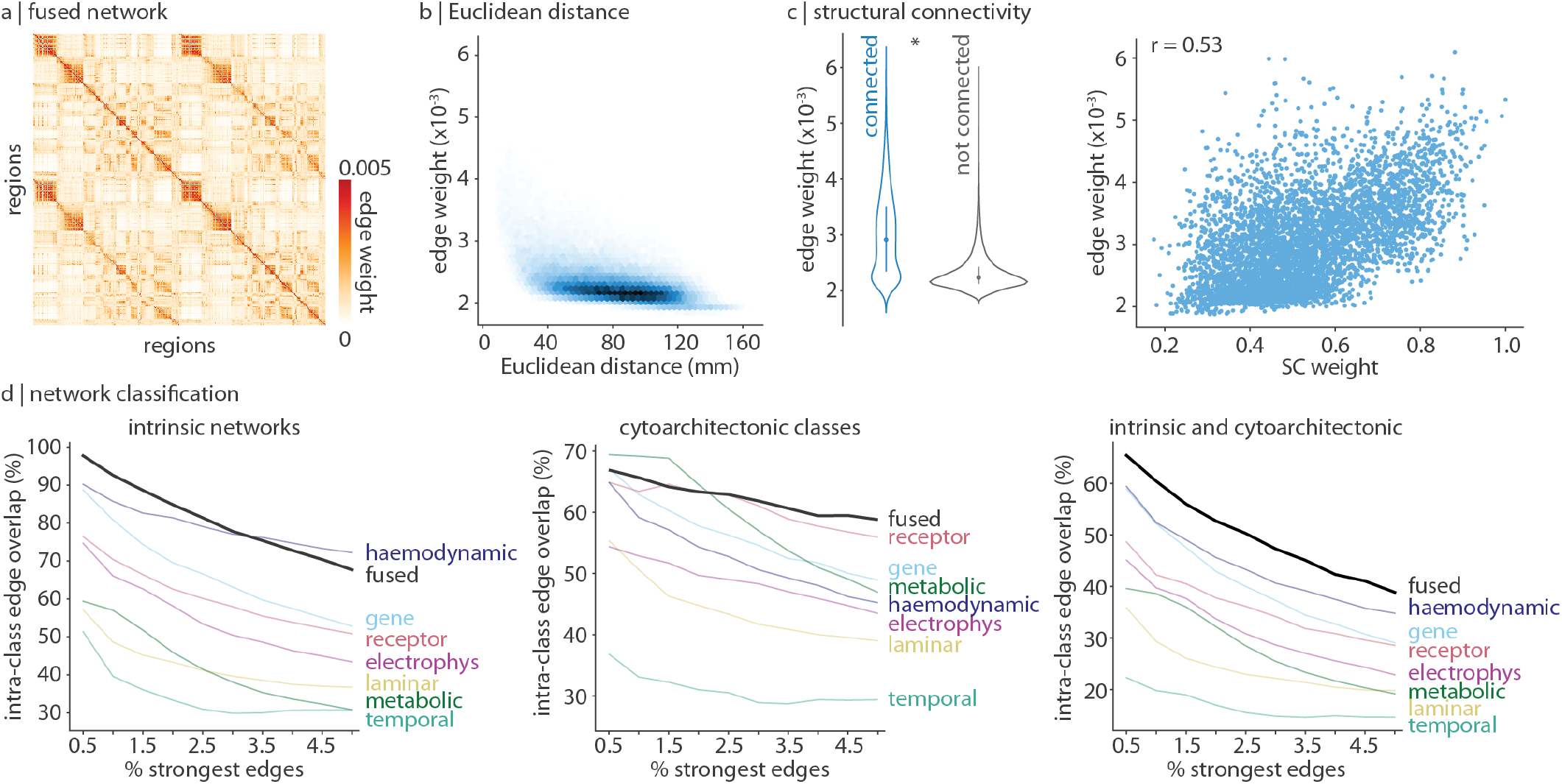
Network fusion. Similarity network fusion was applied to all seven connectivity modes to construct a single integrated network [89, 155]. (a) The fused network. (b) Edge weight decreases exponentially with Euclidean distance. (c) Structurally connected edges have greater edge weight than edges without an underlying structural connection, against a degree and edge-length preserving null model (left [20]), and is correlated with structural connectivity (right). (d) For a varying threshold of strongest edges (0.5%–5% in 0.5% intervals), we calculate the proportion of edges that connect two regions within the same intrinsic network (left), cytoarchitectonic class (middle), and the union of intrinsic networks and cytoarchitectonic classes (right).

We next asked whether the strongest edges in the fused network exist between functionally and cytoarchitectonically similar brain regions (Fig. 6d). We find that nearly all (97.7%) of the top 0.5% strongest edges in the fused network are between regions within the same functional network. Likewise, for cytoarchitectonic classes, we find that the fused network retains more intra-class edges than any other network when the number of strongest edges considered is ≥ 2.5%. Since the fused network represents an integrated cross-modal connectivity mode, we asked whether the strongest edges of the fused network might simultaneously maximize intrinsic and cytoarchitectonic intra-class edges. Indeed, when considering the top 0.5% to 5.0% strongest edges, the number of edges that exist between regions in the same intrinsic and cytoarchitectonic classes is consistently greatest for the fused network. In other words, this multi-modal integrated network potentially recapitulates fundamental organizational properties of the brain better than any isolated network.

### Sensitivity and replication analysis

Finally, to ensure results are not dependent on the parcellation, we repeated all analyses (except Fig. 4 which depends on the 68-region Desikan-Killiany parcellation) using the 100-region Schaefer parcellation and the 68-region Desikan-Killiany parcellation [29, 38, 120]. We find similar results under these alternative parcellations (Fig. S3). These coarser resolutions reveal dense frontal inter-connectivity in the metabolic network, which was not visible at the 400-node parcellation likely due to smoothing effects in dynamic PET data. Furthermore, we share all seven connectivity modes at these three parcellations (Schaefer-400, Schaefer-100, Desikan-Killany-68) in hopes of facilitating integrative connectome analyses in the future (https://github.com/netneurolab/hansen_many_networks).

## DISCUSSION

This work integrates multiple representations of brain connectivity to establish how diverse connectivity modes contribute to brain structure and function. We systematically document the common organizational principles of connectivity modes, as well as their unique contributions to brain structure and geometry. We find that molecular connectivity modes amplify disease exposure resulting in spatial patterns of cortical abnormality. We show that connectivity modes demonstrate diverse dominant gradients and modular structure. Finally, we derive a multimodal, multiscale network by parsimoniously integrating multiple connectivity modes.

Connectomics—the study of relationships between neural elements across multiple scales—is increasingly becoming the dominant paradigm in neuroscience [13, 81, 136]. Numerous technological and analytic methods have been developed to reconstruct inter-regional relationships, some focused on physical wiring, others on molecular similarity, and others still on coherence between regional neural activity. Despite being rooted in common questions, these connectivity modes are often studied in separate literatures. What network features are unique or common to each connectivity mode remains unknown and the practice of studying connectivity modes separately has precluded a truly integrated understanding of inter-regional relationships.

Large-scale comprehensive datasets alongside better data sharing practices have made multi-modal, integrative approaches to studying human brain connectivity more feasible [41, 80, 87, 154]. Examples include comparisons of dynamic FDG-PET and BOLD connectivity [70, 152], BOLD and MEG connectivity [24, 25, 126], structural and BOLD connectivity [65, 144], and correlated gene expression and structural connectivity [113]. Combining connectivity modes has also been used to better resolve clusters of functional activation in BOLD data [84], and inform the application of deep brain stimulation to psychiatric and neurological diseases [11, 67]. Encouragingly, previous work has found that incorporating multiple perspectives of brain connectivity can result in novel discoveries, including improved generative models of brain connectivity [100], structure-function coupling [56, 105], epicentres of transdiagnostic alterations [57, 60], and the characterization of homophilic wiring principles [16].

The present report makes it possible to simultaneously assess relationships between diverse forms of connectivity. Previous literature has repeatedly but separately shown a tendency for homologous [112], spatially proximal [45], and structurally connected [62] brain regions to be similar. In other words, inter-regional relationships appear to be closely related to both geometry and structure [102]. Here we confirm—across multiple connectivity modes, acquired using different imaging modalities, at different spatial scales, and of different biological mechanisms—that these properties are fundamental to brain organization, providing some guidelines with which future connectivity modes can be compared.

However, connectivity modes are not simply redundant. Connectivity modes are poorly correlated among each other. These differences in connection patterns manifest as different principal gradients and modular structure. Even identification of brain hubs—regions that are disproportionately well connected in the network— varies from mode to mode. This suggests that each form of connectivity profile provides a fundamentally different but important view of how regions participate in neural circuits at different spatial and temporal scales [15]. For example, microscale connectivity modes (e.g. correlated gene expression, receptor similarity) are well delineated by a partition based on cytoarchitectonic classes whereas dynamic connectivity modes (e.g. haemodynamic connectivity, electrophysiological connectivity) fit into intrinsic cognitive systems.

In an effort to understand which brain regions are consistently central across many levels of description, we also identify a set of cross-modal hubs. Brain hubs are typically defined as regions with a relatively large number of structural connections. Meanwhile, cross-modal hubs—regions that are consistently similar to the rest of the brain, no matter the connectivity mode—exist in the precuneus, supramarginal gyrus, and dorsolateral prefrontal cortex. These are association regions that most expand during evolution and are involved in high-level cognition including language, planning, and complex executive functions [63, 158], suggesting that these functions are supported by integration across multiple biological scales. This reinforces a well known concept in network science that hub function can be probed and classified using more than just the number of connections, but also how those connections are arranged (e.g., “connector” vs “provincial” hubs) [135]. This work provides a next step for how hubs should be classified based on their biology [5].

We further find a dichotomy between molecular (e.g. correlated gene expression, receptor similarity, laminar similarity) and dynamic (e.g. haemodynamic and electrophysiological connectivity) modes, where molecular modes tend to better support global communication in the brain. For example, we find that molecular feature similarity is significantly increased for links between regions of the brain’s rich club: high-degree regions that show dense inter-connectivity which is thought to improve global communication efficiency and integration [145]. A transcriptional signature of rich club connectivity was previously shown to be driven by genes involved in metabolism, supporting the theory that the brain’s rich club is energetically expensive [27, 35, 47]. Interestingly, we find that metabolic connectivity is increased in rich links, suggesting that the rich club is also synchronized in its energy consumption. The primacy of microscopic features is perhaps unsurprising given that they are thought to support the emergence of macroscopic and dynamic inter-regional interactions.

Molecular feature similarity—particularly correlated gene expression and receptor similarity—also best explain the spatial patterning of cortical disease abnormalities. Recent work has explored the idea that multiple pathologies spread trans-synaptically, including misfolded proteins, aberrant neurodevelopmental signals, and excitotoxic electrical discharge, resulting in patterns of pathology that reflect the underlying structural architecture of the brain [129, 161]. Here we consider the possibility that shared vulnerability is not only limited to structural connections but extends to multiscale biological attributes [57]. We use changes in cortical thickness as the marker of potential pathology and find that when disease exposure is informed by transcriptional and receptor similarity, we can reproduce the cortical profile of multiple diseases (*r >* 0.5 for most). Collectively, molecular similarity is an informative mode of brain connectivity that provides a more comprehensive account for how brain regions may engage in signal exchange and molecular transport.

The present work should be considered alongside some methodological considerations. First, the results are only representative of the seven included connectivity modes; future work should extend this work into additional forms of connectivity. One exciting avenue would be to annotate structural connectomes with measures of myelin or axon caliber derived from quantitative MRI such as magnetization transfer (MT), T1 relaxation rate (R1), or axon diameter [7, 83]. Second, each connectivity matrix is dependent on the quality of the imaging modality, and each imaging method operates at a unique spatial and temporal resolution. Results may therefore be influenced by differences in how the data are acquired. We tried to mitigate this by running extensive sensitivity analyses. Third, in an effort to make correlated gene expression comparable to the other modes, data interpolation and mirroring was conducted, potentially biasing this network towards homotopic connections. Fourth, connectivity modes are compiled across different individuals of varying ages and sex ratios. Results are therefore limited to group-averages, and motivate future deep phenotyping studies of the brain across multiple scales and modalities.

Altogether, this work harmonizes seven perspectives of brain connectivity from diverse spatial scales and imaging modalities including gene expression, receptor density, cellular composition, metabolic consumption, haemodynamic activity, electrophysiology, and time-series features. We demonstrate both the similar and complementary ways in which connectvitiy modes reflect brain geometry, structure, and disease. This serves as a step towards the next-generation integrative, multi-modal study of brain connectivity.

## METHODS

### Connectivity modes

We construct cortical connectivity modes for seven different brain features: gene expression, receptor density, lamination, glucose uptake, haemodynamic activity, electrophysiological activity, and temporal profiles. Each connectivity mode is defined across 400 cortical brain regions, ordered according to 7 intrinsic networks (visual, somatomotor, dorsal attention, ventral attention, limbic, frontoparietal, default mode), separated by hemispheres (left, right) [120]. Replication analyses were conducted using the 100-region Schaefer and 68-region Desikan-Killiany parcellations (all available at https://github.com/netneurolab/hansen_many_networks). To facilitate comparison between networks, each network is normalized using Fisher’s *r*-to-*z* transform (*z* = arctanh(*r*)). We describe the construction of each connectivity mode in detail below.

#### Correlated gene expression

Correlated gene expression represents the transcriptional similarity between pairs of brain regions. Regional microarry expression data were obtained from 6 postmortem brains (1 female, ages 24.0–57.0, 42.50 ± 13.38) provided by the Allen Human Brain Atlas (AHBA, https://human.brain-map.org [59]). Data were processed with the abagen toolbox (version 0.1.1; https://github.com/rmarkello/abagen [86]) using a 400-region volumetric atlas in MNI space.

First, microarray probes were reannotated using data provided by Arnatkevičiūtė et al. [6]; probes not matched to a valid Entrez ID were discarded. Next, probes were filtered based on their expression intensity relative to background noise [107], such that probes with intensity less than the background in ≥ 50% of samples across donors were discarded, yielding 31 569 probes. When multiple probes indexed the expression of the same gene, we selected and used the probe with the most consistent pattern of regional variation across donors (i.e., differential stability [58]), calculated with:

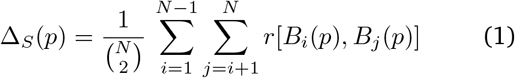

where *ρ* is Spearman’s rank correlation of the expression of a single probe, *p*, across regions in two donors *B*_*i*_ and *B*_*j*_, and *N* is the total number of donors. Here, regions correspond to the structural designations provided in the ontology from the AHBA.

The MNI coordinates of tissue samples were updated to those generated via non-linear registration using the Advanced Normalization Tools (ANTs; https://github.com/chrisfilo/alleninf). To increase spatial coverage, tissue samples were mirrored bilaterally across the left and right hemispheres [113]. Samples were assigned to brain regions in the provided atlas if their MNI coordinates were within 2mm of a given parcel. If a brain region was not assigned a tissue sample based on the above procedure, every voxel in the region was mapped to the nearest tissue sample from the donor in order to generate a dense, interpolated expression map. The average of these expression values was taken across all voxels in the region, weighted by the distance between each voxel and the sample mapped to it, in order to obtain an estimate of the parcellated expression values for the missing region. All tissue samples not assigned to a brain region in the provided atlas were discarded.

Inter-subject variation was addressed by normalizing tissue sample expression values across genes using a robust sigmoid function [49]:

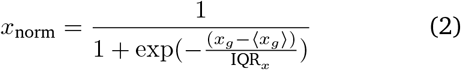

where ⟨*x*⟩ is the median and IQR_*x*_ is the normalized interquartile range of the expression of a single tissue sample across genes. Normalized expression values were then rescaled to the unit interval:

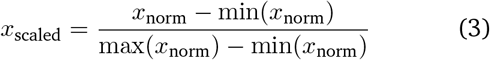

Gene expression values were then normalized across tissue samples using an identical procedure. Samples assigned to the same brain region were averaged separately for each donor, yielding a regional expression matrix for each donor with 400 rows, corresponding to brain regions, and 15 633 columns, corresponding to the retained genes. A threshold of 0.1 was imposed on the differential stability of each gene, such that only stable genes were retained for future analysis, resulting in 8 687 retained genes.

Finally, the region × region correlated gene expression matrix was constructed by correlating (Pearson’s *r*) the normalized gene expression profile at every pair of brain regions. This matrix was then normalized using Fisher’s *r*-to-*z* transform.

#### Receptor similarity

Receptor similarity indexes the degree to which the receptor density profiles at two brain regions are correlated. Conceptually, it can be thought of as how similarly two brain regions might “hear” the same neural signal. PET tracer images for 18 neurotransmitter receptors and transporters were obtained from Hansen et al. [56] and neuromaps (v0.0.1, https://github.com/netneurolab/neuromaps [87]). The receptors/transporters span 9 neurotransmitter systems including: dopamine (D1, D2, DAT), norepinephrine (NET), serotonin (5-HT1A, 5-HT1B, 5-HT2, 5-HT4, 5-HT6, 5-HTT), acetylcholine (*α*_4_*β*_2_, M1, VAChT), glutamate (mGluR5), GABA (GABAA), histamine (H3), cannabinoid (CB1), and opioid (MOR). Tracer names and number of participants (with number of females in parentheses) are listed for each receptor in Table S1. Each PET tracer image was parcellated to 400 brain regions and z-scored. A region-by-region receptor similarity matrix was constructed by correlating (Pearson’s *r*) receptor profiles at every pair of brain regions. This matrix was then normalized using Fisher’s *r*-to-*z* transform.

#### Laminar similarity

Laminar similarity is estimated from histological data and aims to uncover how similar pairs of brain regions are in terms of cellular distributions across the cortical laminae. Specifically, we use data from the Big-Brain, a high-resolution (20 *μ*m) histological reconstruction of a post-mortem brain from a 65 year old male [4, 103]. Cell-staining intensity profiles were sampled across 50 equivolumetric surfaces from the pial surface to the white mater surface to estimate laminar variation in neuronal density and soma size. Intensity profiles at various cortical depths can be used to approximately identify boundaries of cortical layers that separate supragranular (cortical layers I–III) granular (cortical layer IV), and infragranular (cortical layers V-VI) layers.

The data were obtained on *fsaverage* surface (164k vertices) from the BigBrainWarp toolbox [104] and were parcellated into 400 cortical regions according to the Schaefer-400 atlas [120]. The region × region laminar similarity matrix was calculated as the partial correlation (Pearson’s *r*) of cell intensities between pairs of brain regions, after correcting for the mean intensity across brain regions. Laminar similarity was first introduced in Paquola et al. [103]. This matrix was then normalized using Fisher’s *r*-to-*z* transform.

#### Metabolic connectivity

Metabolic connectivity indexes how similarly two brain regions metabolize glucose over time and therefore how similarly two brain regions consume energy. Volumetric 4D PET images of [F^18^]-fluordoxyglucose (FDG, a glucose analogue) tracer uptake over time were obtained from Jamadar et al. [69]. Specifically, 26 healthy participants (77% female, 18–23 years old) were recruited from the general population and underwent a 95 minute simultaneous MR-PET scan in a Siemens (Erlangen) Biograph 3-Tesla molecular MR scanner. Participants were positioned supine in the scanner bore with their head in a 16-channel radiofrequency head coil and were instructed to lie as still as possible with eyes open and think of nothing in particular. FDG (average dose 233 MBq) was infused over the course of the scan at a rate of 36 mL/h using a BodyGuard 323 MR-compatible infusion pump (Caesarea Medical Electronics, Caesarea, Israel). Infusion onset was locked to the onset of the PET scan. This data has been validated and analyzed previously in [70, 152].

PET images were reconstructed and preprocessed according to [152]. Specifically, the 5700-second PET time-series for each subject was binned into 356 3D sinogram frames each of 16-second intervals. The attenuation for all required data was corrected via the pseudo-CT method [28]. Ordinary Poisson-Ordered Subset Expectation Maximization algorithm (3 iterations, 21 subsets) with point spread function correction was used to reconstruct 3D volumes from the sinogram frames. The reconstructed DICOM slices were converted to NIFTI format with size 344 × 344 × 127 (voxel size: 2.09 2.09 2.03 mm^3^) for each volume. A 5 mm FWHM Gaussian postfilter was applied to each 3D volume. All 3D volumes were temporally concatenated to form a 4D (344 × 344 × 127 × 356) NIFTI volume. A guided motion correction method using simultaneously acquired MRI was applied to correct the motion during the PET scan. 225 16-second volumes were retained commencing for further analyses.

Next, the 225 PET volumes were motion corrected (FSL MCFLIRT [71]) and the mean PET image was brain extracted and used to mask the 4D data. The fPET data were further processed using a spatiotemporal gradient filter to remove the accumulating effect of the radiotracer and other low-frequency components of the signal [69]. Finally, each time point of the PET volumetric time-series were registered to MNI152 template space using Advanced Normalization Tools in Python (ANTSpy, https://github.com/ANTsX/ANTsPy), parcellated to 400 regions according to the Schaefer atlas, and time-series at pairs of brain regions were correlated (Pearson’s *r*) to construct a metabolic connectivity matrix for each subject. A group-averaged metabolic connectome was obtained by averaging connectivity across subjects, and lastly the matrix was normalized using Fisher’s *r*-to-*z* transform.

#### Haemodynamic connectivity

Haemodynamic connectivity, commonly simply referred to as “functional connectivity”, captures how similarly pairs of brain regions exhibit fMRI BOLD activity at rest. The fMRI BOLD time-series picks up on magnetic differences between oxygenated and deoxygenated haemoglobin to measure the haemodynamic response: the oversupply of oxygen to active brain regions [76]. Functional magnetic resonance imaging (MRI) data were obtained for 326 unrelated participants (age range 22— 35 years, 145 males) from the Human Connectome Project (HCP; S900 release [147]). All four resting state fMRI scans (two scans (R/L and L/R phase encoding directions) on day 1 and two scans (R/L and L/R phase encoding directions) on day 2, each about 15 min long; TR=720 ms) were available for all participants. Functional MRI data were pre-processed using HCP minimal pre-processing pipelines [53, 147]. Specifically, all 3T functional MRI time-series were corrected for gradient nonlinearity, head motion using a rigid body transformation, and geometric distortions using scan pairs with opposite phase encoding directions (R/L, L/R) [36]. Further pre-processing steps include co-registration of the corrected images to the T1w structural MR images, brain extraction, normalization of whole brain intensity, highpass filtering (>2000s FWHM; to correct for scanner drifts), and removing additional noise using the ICA-FIX process [36, 114]. The pre-processed time-series were then parcellated to 400 cortical brain regions according to the Schaefer atlas [120]. The parcellated time-series were used to construct functional connectivity matrices as a Pearson correlation coefficient between pairs of regional time-series for each of the four scans of each participant. A group-average functional connectivity matrix was constructed as the mean functional connectivity across all individuals and scans. This matrix was then normalized using Fisher’s *r*-to-*z* transform.

#### Electrophysiological connectivity

Electrophysiological connectivity was measured using magnetoencephalography (MEG) recordings, which tracks the magnetic field produced by neural currents. Resting state MEG data was acquired for *n* = 33 unrelated healthy young adults (age range 22–35 years) from the Human Connectome Project (S900 release [147]). The data includes resting state scans of approximately 6 minutes long and noise recording for all participants. MEG anatomical data and 3T structural MRI of all participants were also obtained for MEG pre-processing. After preprocessing and parcellating the data, amplitude envelope correlations were performed between time-series at each pair of brain regions, for six canonical frequency bands separately (delta (2–4 Hz), theta (5–7 Hz), alpha (8–12 Hz), beta (15–29 Hz), low gamma (30–59 Hz), and high gamma (60–90 Hz)). Amplitude envelope correlation is applied instead of directly correlating the time-series because of the high sampling rate (2034.5 Hz) of the MEG recordings. The composite electrophysiological connectivity matrix is the first principal component of all six connectivity matrices (vectorized upper triangle), and closely resembles alpha connectivity (Fig. S5). Finally, the matrix underwent Fisher’s *r*-to-*z* transform. More processing details are described below.

The present MEG data was first processed and used by Shafiei et al. [126]. Resting state MEG data was preprocessed using the open-source software, Brainstorm (https://neuroimage.usc.edu/brainstorm/ [140]), following the online tutorial for the HCP dataset (https://neuroimage.usc.edu/brainstorm/Tutorials/HCP-MEG). MEG recordings were registered to individual structural MRI images before applying the following preprocessing steps. First, notch filters were applied at 60, 120, 180, 240, and 300 Hz, followed by a high-pass filter at 0.3 Hz to remove slow-wave and DC-offset artifacts. Next, bad channels from artifacts (including heartbeats, eye blinks, saccades, muscle movements, and noisy segments) were removed using Signal-Space Projections (SSP).

Pre-processed sensory-level data was used to construct a source estimation on HCP’s fsLR4k cortex surface for each participant. Head models were computed using overlapping spheres and data and noise covariance matrices were estimated from resting state MEG and noise recordings. Linearly constrained minimum variance (LCMV) beamformers was used to obtain the source activity for each participant. Data covariance regularization was performed and the estimated source variance was normalized by the noise covariance matrix to reduce the effect of variable source depth. All eigenvalues smaller than the median eigenvalue of the data covariance matrix were replaced by the median. This helps avoid instability of data covariance inversion caused by the smallest eigenvalues and regularizes the data covariance matrix. Source orientations were constrained to be normal to the cortical surface at each of the 8 000 vertex locations on the cortical surface, then parcellated according to the Schaefer-400 atlas [120]. Parcellated time-series were then used to estimate amplitude-based connectivity [26]. An orthogonalization process was applied to correct for the spatial leakage effect by removing all shared zero-lag signals [33].

#### Temporal profile similarity

Temporal profile similarity was first introduced by, and obtained from, Shafiei et al. [128] and represents how much two brain regions exhibit similar temporal features, as calculated on fMRI time-series. Specifically, we used the highly comparative time-series analysis toolbox, *hctsa* [48, 49] *to perform a massive feature extraction of the parcellated fMRI time-series (see Haemodynamic connectivity*) at each brain region of each participant. The *hctsa* package extracted over 7 000 local time-series features using a wide range of operations based on time-series analysis. The extracted features include, but are not limited to, distributional features, entropy and variability, autocorrelation, time-delay embeddings, and nonlinear features of a given time-series. Following the feature extraction procedure, the outputs of the operations that produced errors were removed and the remaining features (6 441 features) were normalized across nodes using an outlier-robust sigmoidal transform. We used Pearson’s correlation coefficients to measure the pairwise similarity between the time-series features of all possible combinations of brain areas. As a result, a temporal profile similarity network was constructed for each individual and each run, representing the strength of the similarity of the local temporal fingerprints of brain areas. This matrix was then normalized using Fisher’s *r*-to-*z* transform.

### Structural connectivity

Diffusion weighted imaging (DWI) data were obtained for 326 unrelated participants (age range 22-35 years, 145 males) from the Human Connectome Project (HCP; S900 release [147]) [36]. DWI data was pre-processed using the MRtrix3 package [143] (https://www.mrtrix.org/). More specifically, fiber orientation distributions were generated using the multi-shell multi-tissue constrained spherical deconvolution algorithm from MRtrix [39, 72]. White matter edges were then reconstructed using probabilistic streamline tractography based on the generated fiber orientation distributions [142]. The tract weights were then optimized by estimating an appropriate cross-section multiplier for each streamline following the procedure proposed by Smith et al. [133] and a connectivity matrix was built for each participant using the 400-region Schaefer parcellation [120]. A group-consensus binary network was constructed using a method that preserves the density and edge-length distributions of the individual connectomes [21, 92, 93]. Edges in the group-consensus network were assigned weights by averaging the log-transformed streamline count of non-zero edges across participants. Edge weights were then scaled to values between 0 and 1.

### Disease exposure

Patterns of cortical thickness from the ENIGMA consortium and the *enigma* toolbox were available for thirteen neurological, neurodevelopmental, and psychiatric disorders (https://github.com/MICA-MNI/ENIGMA; [57, 79, 141]), including: 22q11.2 deletion syndrome (*N* = 474 participants, *N* = 315 controls) [138], attention-deficit/hyperactivity disorder (ADHD; *N* = 733 participants, *N* = 539 controls) [66], autism spectrum disorder (ASD; *N* = 1571 participants, *N* = 1651 controls) [148], idiopathic generalized (*N* = 367 participants), right temporal lobe (*N* = 339 participants), and left temporal lobe (*N* = 415 participants) epilepsies (*N* = 1727 controls) [157], depression (*N* = 2148 participants, *N* = 7957 controls) [121], obsessive-compulsive disorder (OCD; *N* = 1905 participants, *N* = 1760 controls) [23], schizophrenia (*N* = 4474 participants, *N* = 5098 controls) [146], bipolar disorder (*N* = 1837 participants, *N* = 2582 controls) [61], obesity (*N* = 1223 participants, *N* = 2917 controls) [101], schizotypy (*N* = 3004 participants) [75], and Parkinson’s disease (*N* = 2367 participants, *N* = 1183 controls) [77]. The ENIGMA (Enhancing Neuroimaging Genetics through Meta-Analysis) Consortium is a data-sharing initiative that relies on standardized processing and analysis pipelines, such that disorder maps are comparable [141]. Altogether, over 21 000 participants were scanned across the thirteen disorders, against almost 26 000 controls. The analysis was limited to adults in all cases except ASD where the cortical abnormality map is only available aggregated across all ages (2–64 years). The values for each map are z-scored effect sizes (Cohen’s *d*) of cortical thickness in patient populations versus healthy controls. Imaging and processing protocols can be found at http://enigma.ini.usc.edu/protocols/. Local review boards and ethics committees approved each individual study separately, and written informed consent was provided according to local requirements.

We calculate disease exposure for every disease and network, after masking the network such that all edges with negative strength are assigned a strength of 0. For a given network and disease, disease exposure of a node *i* is defined as,

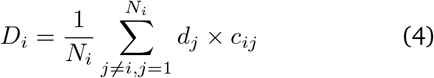

where *N*_*i*_ is the number of positive connections made by region *i, d*_*j*_ is the cortical abnormality at region *j*, and *c*_*ij*_ is the edge strength between regions *i* and *j*. This analysis was repeated after regressing the exponential fit in Fig. 1b from each network, to ensure results are not driven by distance (Fig. S4).

### Community detection

For each connectivity mode, communities were identified using the Louvain algorithm, which maximizes positive edge strength within groups and negative edge strength between groups [22]. Specifically, communities were assigned to nodes in a manner that maximizes the quality function

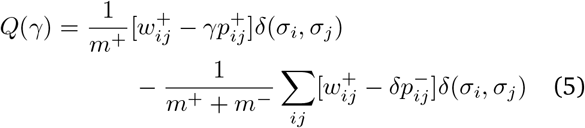

where 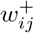 is the network with only positive correlations and likewise for 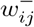 and negative correlations. The term 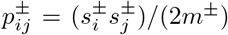 represents the null model: the expected density of connections between nodes *i* and *j*, where 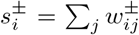 and 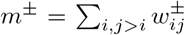. The variable *σ*_*i*_ is the community assignment of node *i* and *δ*(*σ*_*i*_, *σ*_*j*_) is the Kronecker function and is equal to 1 when *σ*_*i*_ = *σ*_*j*_ and 0 otherwise. The resolution parameter, *γ*, scales the relative importance of the null model, making it easier (*γ >* 1) or harder (*γ <* 1) for the algorithm to uncover many communities. In other words, as *γ* increases, increasingly fine network partitions are identified. We tested 60 values of *γ*, from *γ* = 0.1 to *γ* = 6.0, in increments of 0.1. At each *γ*, we repeated the algorithm 250 times and constructed a consensus partition, following the procedure presented in Bassett et al. [12].

### Similarity network fusion

First introduced by Wang et al. [155], similarity network fusion (SNF) is a method for combining multiple measurement types for the same observations (e.g. patients, or in our case, brain regions) into a single similarity network where edges between observations represent their cross-modal similarity. For each data source, SNF constructs an independent similarity network, defines the *K* nearest neighbours for each observation, then iteratively combines the networks in a manner that gives more weight to edges between observations that are consistently high-strength across data types. We relied on snfpy (https://github.com/rmarkello/snfpy [89]), an open-source Python implementation of the original SNF code provied by Wang et al. [155]. A brief description of the main steps in SNF follows, adapted from its original presentation in Wang et al. [155].

In the present report, the seven data sources to be fused are the seven connectivity modes (correlated gene expression, receptor similarity, laminar similarity, metabolic connectivity, haemodynamic connectivity, electrophysiological connectivity, and temporal similarity). First, similarity networks for each connectivity mode is constructed where edges are determined using a scaled exponential similarity kernel:

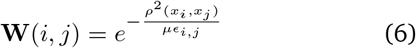

where **W**(*i, j*) is the edge weight between regions *i* and *j, ρ*(*x*_*i*_, *x*_*j*_) is the Euclidean distance between regions *i* and *j, μ* ∈ ℝ is a hyperparameter that is set empirically, and

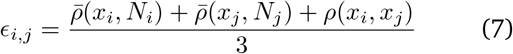

where 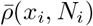 is the average distance between *x*_*i*_ and all other regions in the network. Note that *μ* is a scaling factor that determines the weighting of edges between regions in the similarity network, and is set to *μ* = 0.5 in the present report.

Next, each **W** is normalized such that:

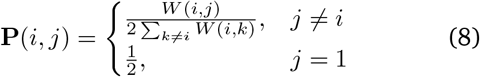

Finally, a sparse matrix **S** of the *K* nearest (i.e. strongest) neighbours is constructed:

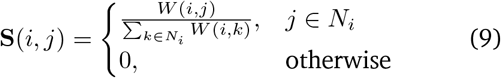

In other words, the matrix **P** encodes the full information about the similarity of each region to all other regions (within a given connectivity mode), whereas **S** encodes only the similarity of the *K* most similar regions to each region. *K* is SNF’s second hyperparameter, which we set to one tenth the number of regions in the network (40).

The similarity networks are then iteratively fused. At each iteration, the matrices are made more similar to each other via:

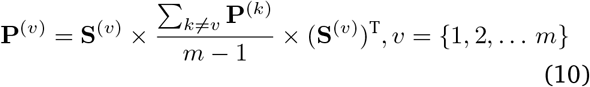

After each iteration, the generated matrices are renormalized as in Equation 8. Fusion stops when the matrices have converged or after a specified number of iterations (in our case, 20). Regions *x*_*i*_ and *x*_*j*_ will likely be neighbours in the fused network if they are neighbours in multiple similarity networks. Furthermore, if *x*_*i*_ and *x*_*j*_ are not very similar in one data type, their similarity can be expressed in another data type.

After the fusion process, we confirm that no single network exerts undue influence on the final fused network by repeating the fusion process while excluding a single network. The minimum correlation (Spearman *r*) between the leave-one-out fused network and the complete fused network is 0.958. In addition to this, we confirm that alternative *K* and *μ* parameters would not make large difference to the fused network. We test *K* ∈ [20, 59] and *μ* ∈ [0.3, 0.8] and find that these alternative fused networks are highly correlated with the original (minimum Spearman *r* = 0.924).

### Null models

*Spin tests* Spatial autocorrelation-preserving permutation tests were used to assess statistical significance of associations across brain regions, termed “spin tests” [3, 88, 149]. We created a surface-based representation of the parcellation on the FreeSurfer fsaverage surface, via files from the Connectome Mapper toolkit (https://github.com/LTS5/cmp). We used the spherical projection of the fsaverage surface to define spatial co-ordinates for each parcel by selecting the coordinates of the vertex closest to the center of the mass of each parcel [150]. These parcel coordinates were then randomly rotated, and original parcels were reassigned the value of the closest rotated parcel (10 000 repetitions). Parcels for which the medial wall was closest were assigned the value of the next most proximal parcel instead. The procedure was performed at the parcel resolution rather than the vertex resolution to avoid upsampling the data, and to each hemisphere separately.

*Network randomization* Structural networks were randomized using a procedure that preserves the density, edge length, degree distributions of the empirical network [20, 149]. Edges were binned according to Euclidean distance (10 bins). Within each bin, pairs of edges were selected at random and swapped, for a total number of swaps equal to the number of regions in the network multiplied by 20. This procedure was repeated 1 000 times to generate 1 000 null structural networks, which were then used to generate null distributions of network-level measures.

## Code and data availability

All code and data used to perform the analyses are available at https://github.com/netneurolab/hansen_many_networks.

## Acknowledgements

We thank Vincent Bazinet, Zhen-Qi Liu, Filip Milisav, Laura Suarez, and Andrea Luppi for their comments and suggestions on the manuscript, the Monash University Neural Systems and Behviour Lab for insightful discussion, and all individuals involved in making the employed open-source datasets available. BM acknowledges support from the Natural Sciences and Engineering Research Council of Canada (NSERC), Canadian Institutes of Health Research (CIHR), Brain Canada Foundation Future Leaders Fund, the Canada Research Chairs Program, the Michael J. Fox Foundation, and the Healthy Brains for Healthy Lives initiative. JYH acknowledges support from the Helmholtz International BigBrain Analytics & Learning Laboratory, the Natural Sciences and Engineering Research Council of Canada, and The Neuro Irv and Helga Cooper Foundation. SDJ acknowledges support from the National Health and Medical Research Council of Australia (APP1174164). The funders had no role in study design, data collection and analysis, decision to publish or preparation of the manuscript.

---

**Figure S1.**
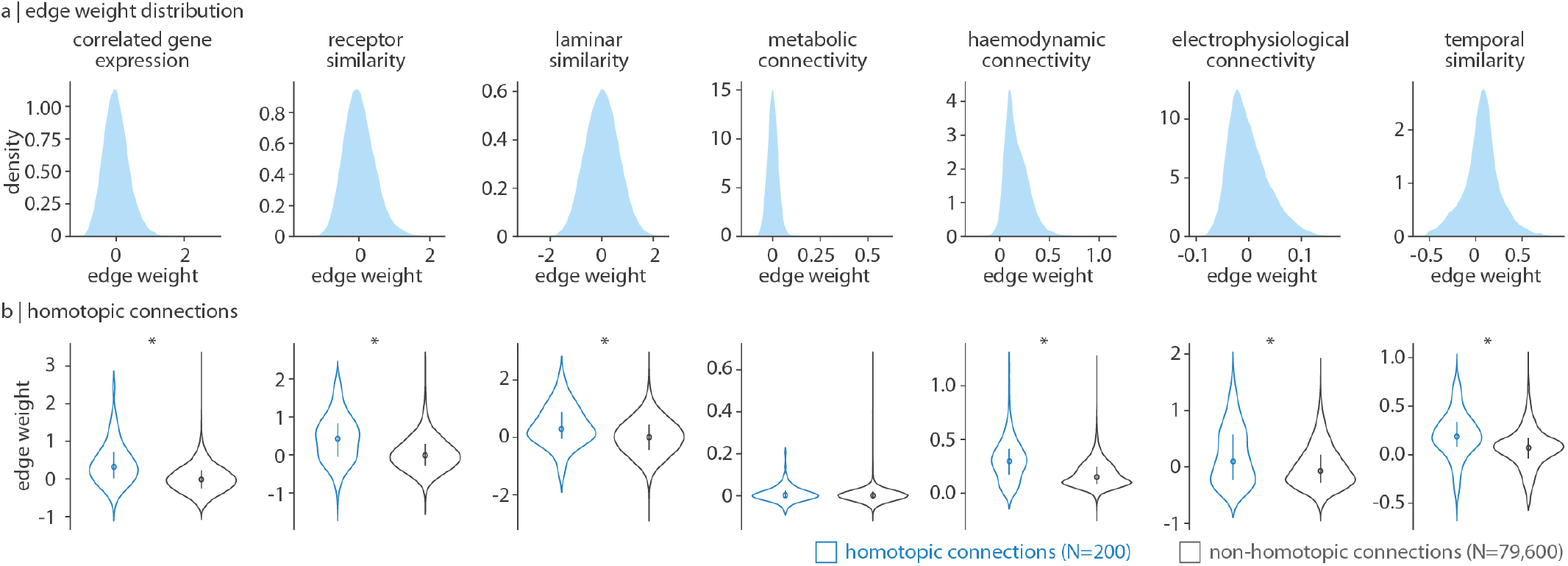
(a) Edge weight distributions of each normalized connectivity mode. (b) Edge weight distributions for homotopic (*N* = 200 edges; blue) and non-homotopic (*N* = 79 600; grey) edges. Significance was determined using two-tailed Welch’s t-test.

**Figure S2.**
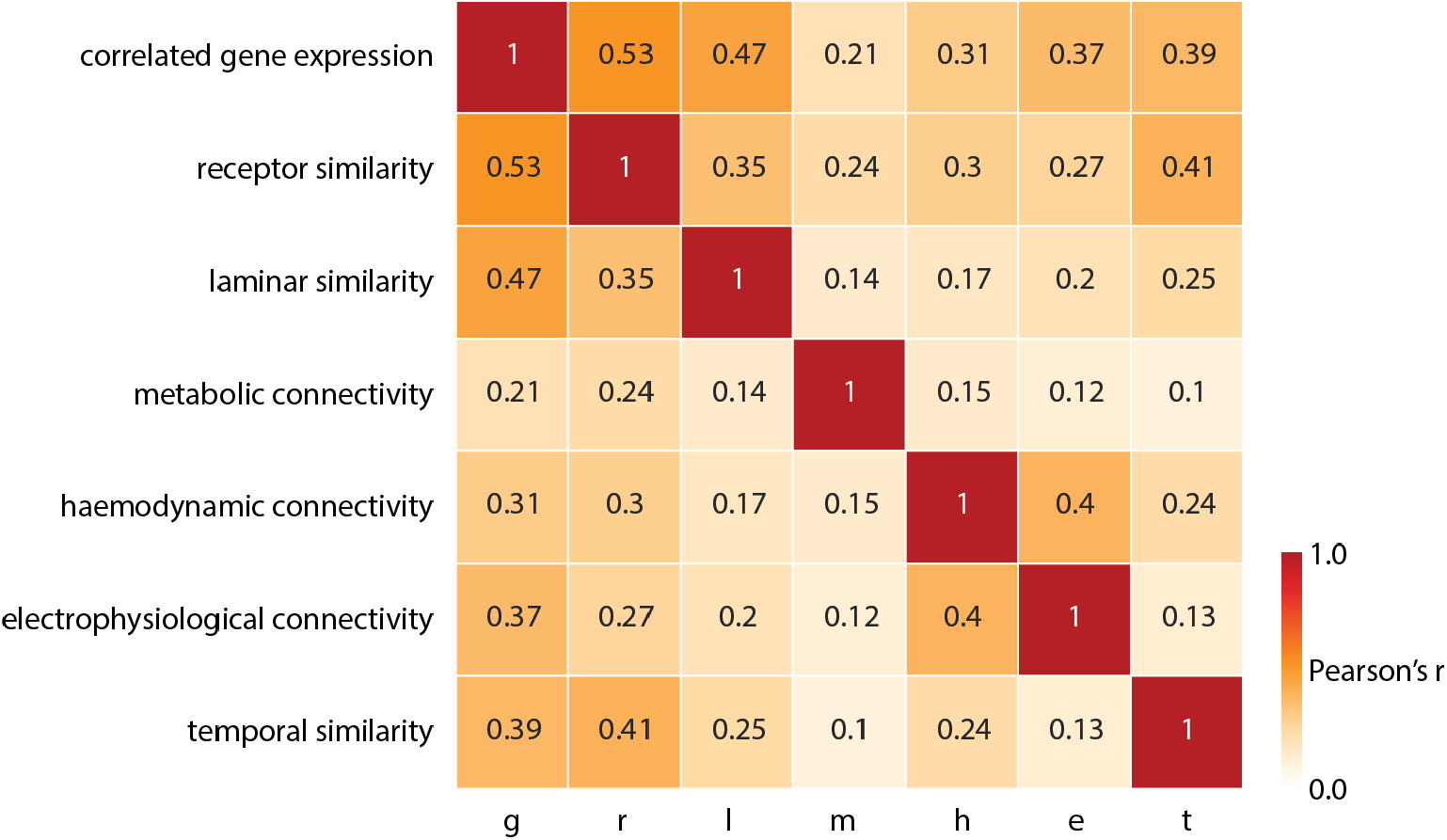
Pearson’s correlation of the upper triangle of pairs of every pair of connectivity modes included in the analysis.

**Figure S3.**
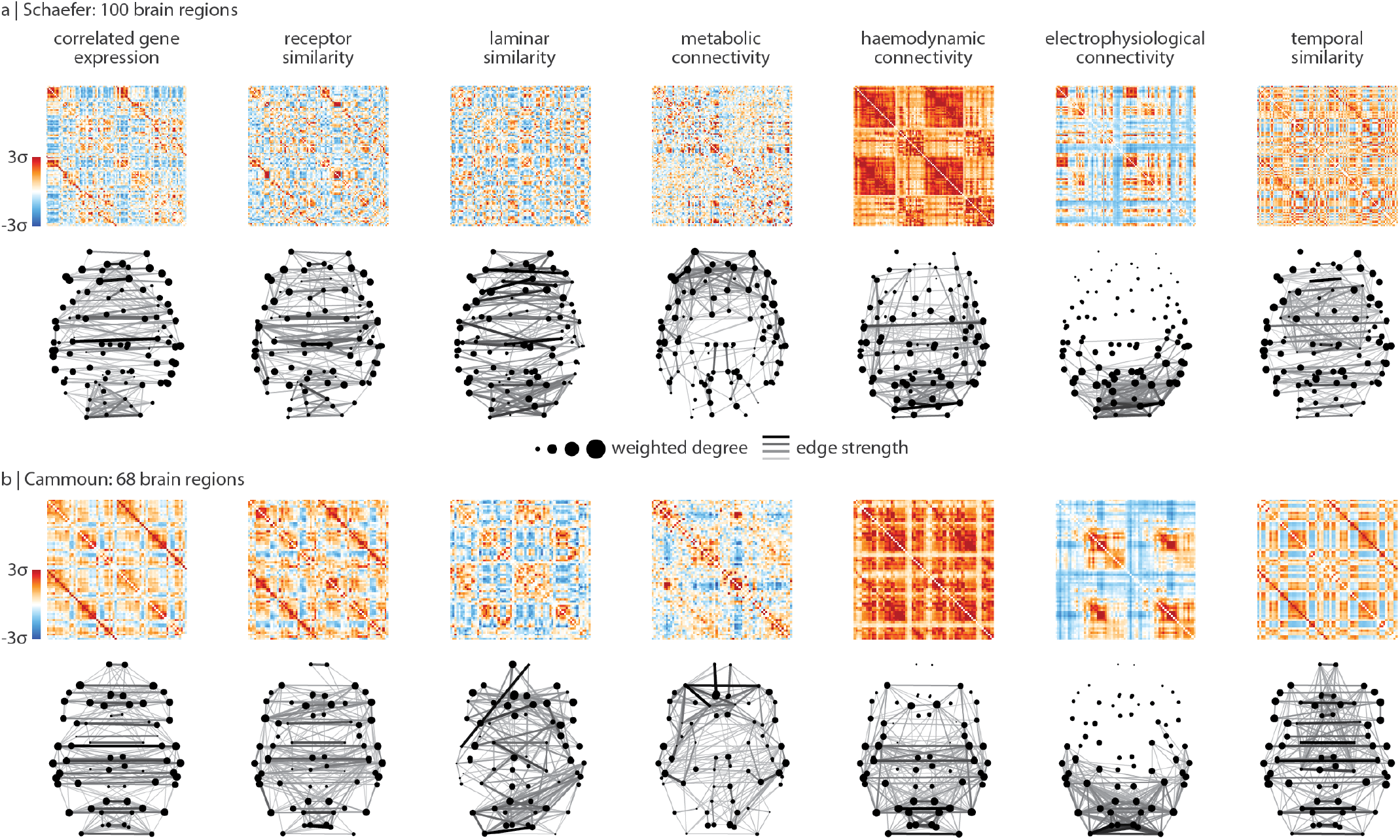
Replication using alternative parcellations. Analyses were repeated using (a) the 100-region Schaefer parcellation and (b) the 68-region Cammoun parcellation (regionally equivalent to the Desikan-Killiany atlas) [29, 38, 120]. We show each network as well as the the 5% (for Schaefer) or 10% (for Cammoun) strongest edges of the network.

**Figure S4.**
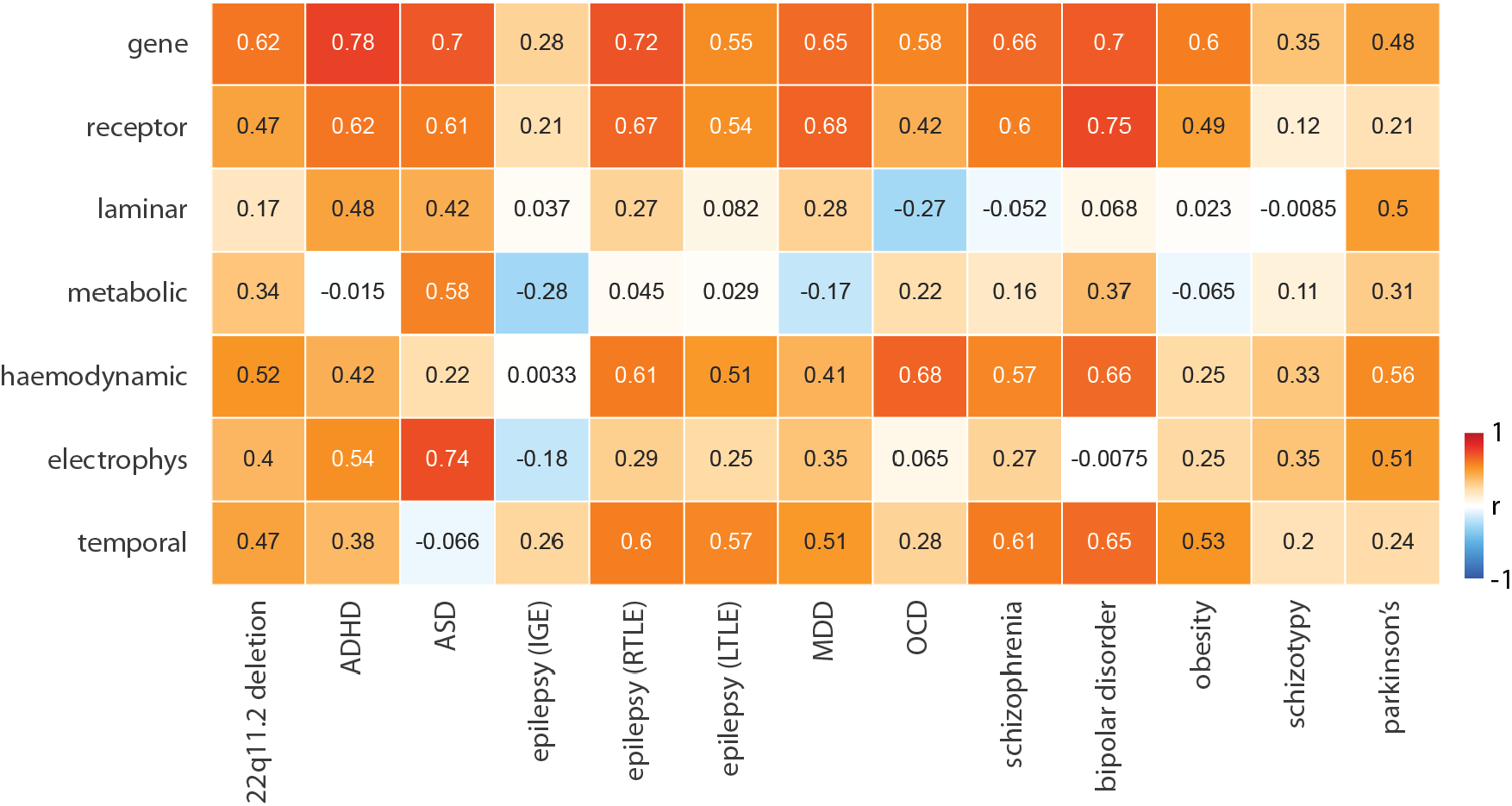
We repeat Fig. 4 after regressing the exponential relationship with distance (shown in Fig. 1b) from each connectivity mode.

**Figure S5.**
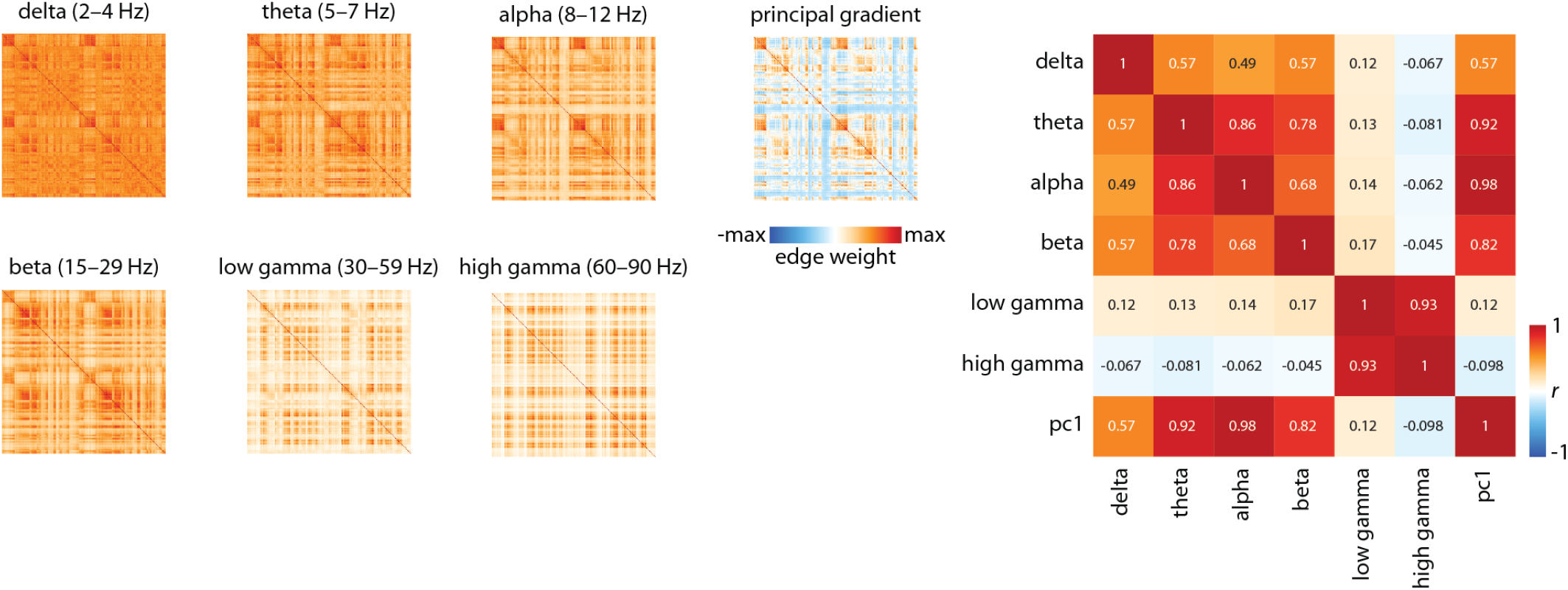
Amplitude envelope correlations were performed between time-series at each pair of brain regions for six canonical frequency bands separately (delta (2–4 Hz), theta (5–7 Hz), alpha (8–12 Hz), beta (15–29 Hz), low gamma (30–59 Hz), high gamma (60–90 Hz). The electrophysiological connectivity mode used in the present analyses is the first principal component of the vectorized upper triangles of all six frequency-dependent connectivity matrices. On the right we show Pearson’s correlations between the six frequency-dependent connectivity matrices and the principal gradient.

**TABLE S1.**
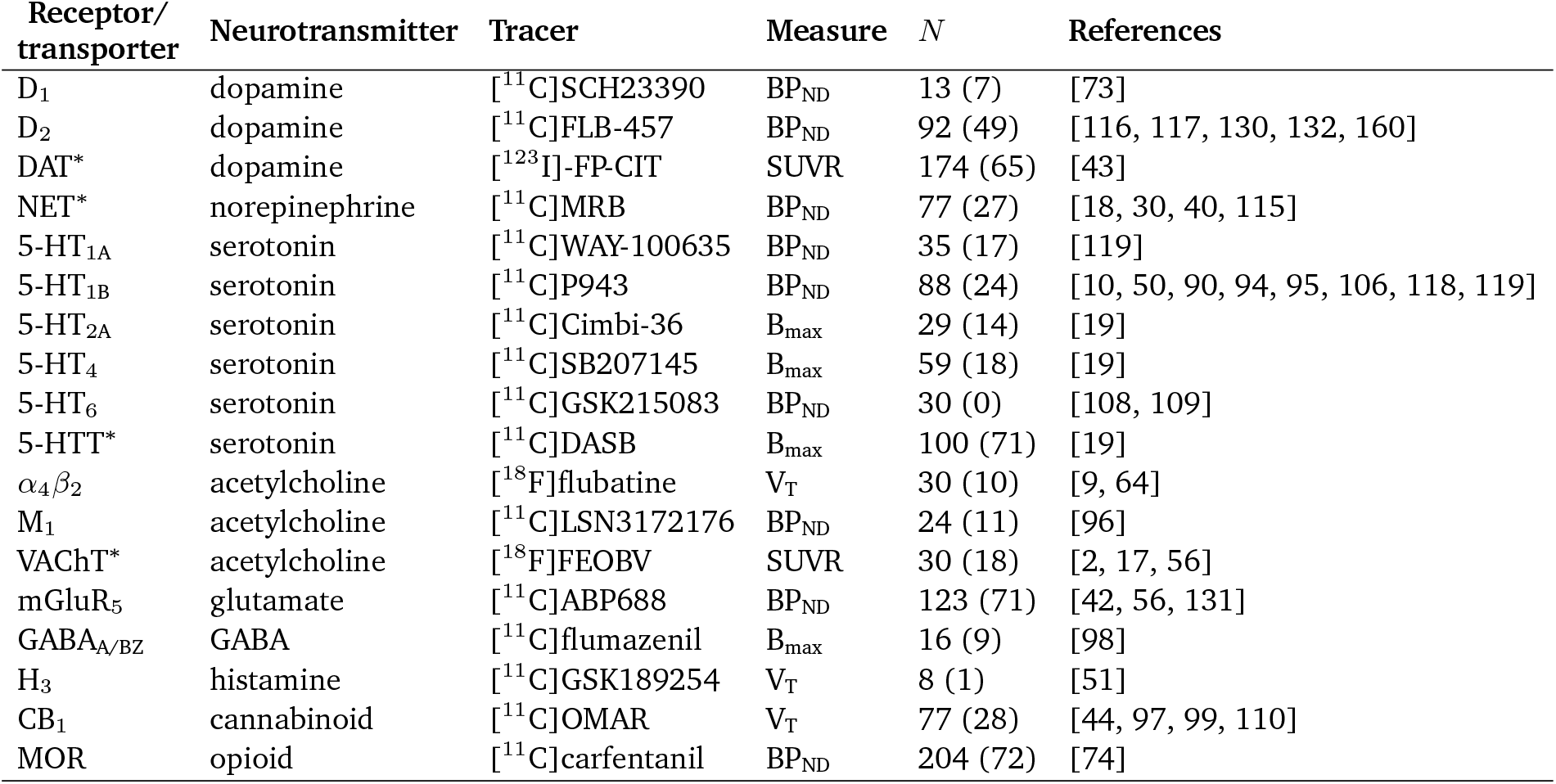
Neurotransmitter receptors and transporters included in receptor similarity. BP_ND_ = non-displaceable binding potential; V_T_ = tracer distribution volume; B_max_ = density (pmol/ml) converted from binding potential (5-HT) or distributional volume (GABA) using autoradiography-derived densities; SUVR = standard uptake value ratio. Values in parentheses (under *N*) indicate number of females. Asterisks indicate transporters. This table is adapted from Table 1 of [56].

